# Separating noise and function in systems of animal communication: a comparative study of aggressive signaling in crayfish

**DOI:** 10.1101/2020.08.03.234419

**Authors:** Zackary A. Graham, Michael J. Angilletta

## Abstract

A primary issue in the study of dishonest signaling is the researcher’s ability to detect and describe a signal as being dishonest. However, by understanding the relative honesty of a signal as a statistical property of an individual or population, researchers have recently quantitively describe dishonest communication. Thus, dishonesty signals can be understood as when there is a breakdown in the correlation between a signal and its underlying meaning; creating variation within a signaling system. However, such variation in signaling systems may not be attributed to dishonesty, because of inherent noise within biological systems driven by evolutionary or physiological noise. Here, we try to separate out functional variation within honest or dishonesty signaling systems from inherent biological noise by leveraging homologous structures that have evolved for separate functions – the enlarged claws of freshwater crayfish. Because burrowing species of freshwater crayfish claws have not evolved as signals, the variability in the size and strength of their claws should be minimal when compared to claws of non-burrowing species that evolved as signals during aggression. We found that despite the claws of burrowing and nonburrowing crayfish claws having evolved to serve difference functions, the claws of all species in our study were inherently noisy. Furthermore, although claws that unreliably correlate to the strengthen the wielder may function as dishonest signals in other crustaceans, we did not find support for this hypothesis; because crayfish escalated aggression based on relative body size.

## INTRODUCTION

Morphological structures that function as signals often vary among individuals more than do other structures. This hypervariability enables receivers to distinguish signalers and to use these structures to observe the welders quality through assessment of size or magnitude of the signal. In many cases, this hypervariability exists because signal size reflects the condition of the signaler (Cotton, Fowler, & Pomiankowski, 2004; Cotton, Small, & Pomiankowski, 2006; Patrice David, Bjorksten, Fowler, & Pomiankowski, 2000). The energy invested in a signaling structure tells receivers the genetic or physical quality of signaler. Typically, an individual of highest quality produces the largest and most extravagant signal (Searcy & Nowicki, 2005). By contrast, structures that evolved for a functions other than signaling (e.g. locomotion, feeding) usually lies in a narrow range of sizes that optimize performance, enabling its bearer to survive and reproduce (Arnold, 1983). Selection for performance traits reduces variation by eliminating alleles that push phenotypes away from to an optimal size. Therefore, signaling structures and performance structures each face unique selection pressures that are predicted to either increase or decrease the variation in the sizes and performance of structures. The eyestalks of *Diopsid* flies exemplify this contrasting variability between signaling traits and performance traits. The eyestalks, which males use to signal their quality to females, vary greatly among individuals of a given body size; by comparison the size of wings varies little among males, as do the non-signaling eyestalks of female flies (David et al., 1998; Patrice David et al., 2000; Panhuis & Wilkinson, 1999). Thus, by comparing the variability between a signal (e.g., male eyestalks) and performance structures (e.g., female eyestalks, wings) biologists can separate the degree of functional variation in signaling systems from outside sources of variation (Graham, Garde, Heide-Jørgensen, & Palaoro, 2020; O’Brien et al., 2018).

Dishonest signaling increases variation within a signaling system, which can ultimately break down the covariation between a trait and the quality being signaled (reviewed in Wilson & Angilletta, 2015). For example, an atypical morph of the fig wasp (*Philotrypesis sp*.) wields exaggerated mandibles that are used as signals during male-male combat (Moore, Obbard, Reuter, West, & Cook, 2009). In most males, the size of the mandible accurately reflects fighting ability, but some males fight poorly despite having disproportionately large mandibles (Moore et al., 2009). The latter males benefit from signaling dishonestly as long as an interactions between males ends before escalating to combat. Selection for dishonest signals weakens the correlation between signals and quality expected in a system of honest signaling. Thus, when selection favors dishonest signaling, we should see greater variation in the quality of individuals with equivalent signals (Elwood & Briffa, 2006; Funk & Tallamy, 2000; Muramatsu & Koga, 2016; Wilson, Angilletta, James, Navas, & Seebacher, 2007). However, the extent of this variation depends on the frequency and intensity of social policing, which imposes a cost of punishment that offsets the expected benefit of deception (Bywater & Wilson, 2012; Catteeuw, Han, & Manderick, 2014; Tibbetts & Izzo, 2010). If social policing is nonexistent, dishonesty will run rampant and promote increased variability in the form of large but poorly quality signals. Ultimately, the benefits of dishonest signaling and the costs of social policing will determine the degree of dishonesty within a communication system (Bywater & Wilson, 2012; Catteeuw et al., 2014; Tibbetts & Izzo, 2010).

Although hypervariability of signaling structures has been interpreted as evidence of dishonest signaling, such variation is necessary but insufficient to establish that a structure serves as a signal. For a given body size, one should expect inherent variation in the size, shape, or performance of any structure to arise from random factors during development or evolution (Eldar & Elowitz, 2010; Richard & Yvert, 2014; Tsimring, 2014; Viney & Reece, 2013). Further, although researchers have recognized the roles that developmental noise and genetic drift play in animal communication (Brumm, 2013; Tanner & Bee, 2019), we need comparative studies to disentangle how much variation stems from selection for optimal performance and how much variation stems from signaling versus other processes. To address this problem, we studied the variation in size and performance of crayfish claws—homologous structures that serve as signals of strength, weapons of aggression, or tools for burrowing. We contrasted phenotypic variation between two groups of species: burrowing species that use claws primarily as tools for digging, and non-burrowing species that use claws primarily as weapons for fighting. Because the claws of non-burrowing crayfish species presumably function as signals during aggression, we expected non-burrowing crayfish to exhibit more variation in the relative size and strength of claws. Furthermore, greater variation in the size and strength of non-burrowing species have been proposed to function as dishonest signals; in which some individuals wield large but weakly performing claws. By contrast, we expected minimal variation in the size and strengths of burrowing crayfish claws; which have presumably evolved as performance structures and face reduced selection for variation. We also expected claw size and strength to influence the outcomes of staged territorial interactions between non-burrowing crayfish. Finally, in non-burrowing but not burrowing species, we predicted greater phenotypic variation among males compared to females, because male crayfish of non-burrowing species are more aggressive and should benefit more from having variable, and potentially dishonesty signaling claws. Ultimately, our study provides a framework for separating inherent noise from functional variation such as dishonest signaling in systems of animal communication.

## METHODS

### Study species

We studied six species of crayfish: three species described as primary burrowers (*Cambarus monongalensis, Cambarus. dubius, Lacunicambus thomai*), and three species described as tertiary burrowers (*Cambarus carinirostris, Cambarus. robustus*, and *Faxonius obscurus*). The docile primary burrowers (hereafter referred to as burrowing crayfish) construct extensive burrows in which they spend most of their life (Hobbs, 1981). The aggressive tertiary burrowers (hereafter referred to as non-burrowing crayfish) construct rudimentary burrows under refuge (Hobbs, 1981). Males and females of all six species possess enlarged frontal claws that perform distinct functions. Because burrowing species lack aggression or territorial behavior (Dalosto, Palaoro, Costa, & Santos, 2013; Guiaşu, Saleh, Mozel, & Dunham, 2005), their claws have been selected for the ability to burrow. By contrast, non-burrowing crayfish possess enlarged claws used as weapons and potentially as signals of strength (Moore, 2007; Robinson & Gifford, 2019; Wilson et al., 2007). In some non-burrowing species, males produce large but weak claws that dishonestly signal their strength to competitors (Robinson & Gifford, 2019; Wilson & Angilletta, 2015). Variation in the size or performance of claws within burrowing species serves as a reference to the degree of variation expected in non-burrowing species without adaptation of claws for signaling. Therefore, these two groups of species enable one to contrast the variation in a trait selected that has been selected as a signal or for digging performance.

### Collection and husbandry of animals

We collected 359 crayfish representing the six species throughout Pennsylvania and West Virginia, U.S.A from June 2019 to August 2019 (Table 1). All non-burrowing species were either captured by hand or net. By contrast, burrowing species construct burrows in mud close to the water table. Thus, burrowing species were caught by flooding burrows with water and plunging the burrow with a closed fist. Usually, the resident crayfish surfaced to investigate the disturbance, enabling one to capture the animal. Often, multiple bouts of plunging were required to capture a burrowing crayfish. Because burrowing species are harder to locate and capture, we collected fewer individuals of burrowing species than we did for non-burrowing species (see Table 1). The sex of each crayfish was identified by the presence or absence of male gonopods.

**Table 1.** Total sample size, number of males, number of females and burrowing status each individual species included in this study. NB= non-burrower, B= burrower.

| Species | N (males, females) | Burrowing status |
| --- | --- | --- |
| *Cambarus carinirostris* | 114 (57,57) | NB |
| *Cambarus robustus* | 61 (30,31) | NB |
| *Faxonius obscurus* | 82 (47,35) | NB |
| *Cambarus monongalensis* | 40 (17,23) | B |
| *Cambarus dubius* | 34 (14,20) | B |
| *Lacunicambus thomai* | 28 (17,11) | B |

Each crayfish was held in plastic container (14 × 9 × 6 cm) with 5 cm of dechlorinated tap water. Prior to any data collection, all crayfish were acclimated to laboratory conditions for two days to diminish effects of prior experience (Bergman et al., 2003; Daws, Konzen, & Moore, 2002; Graham, Padilla-Perez, & Angilletta, 2020). While housed in the laboratory, crayfish were not fed and were kept on a 14:10 light day cycle. After data collection, each crayfish was returned to the exact location of capture.

### Measuring the sizes and strengths of claws

The claws of each crayfish were photographed on background of graph paper. Images were imported to ImageJ (ref), a software package that enabled us to quantify six linear dimensions of each claw: 1) width at heel, 2) width at dactyl/manus joint, 3) length of manus from heel to joint, 4) width of pollex at dactyl joint, 5) width of dactyl, and 6) length of pollex (Fig 1). We converted the six dimensions to principle components, yielding a first principle component (PC1) describing more than 93.92% of the variation. As in previous studies of crayfish ((Bywater, Angilletta, & Wilson, 2008; Graham, Padilla-Perez, et al., 2020; Robbie S Wilson et al., 2007; Robbie S Wilson, James, Bywater, & Seebacher, 2009), the scores of the first principal component were used as a measurement of claw size. To relate claw size to body size, we estimated body size as the length of the cephalothorax measured with digital calipers (Graham, Padilla-Perez, et al., 2020).

**Figure 1.**
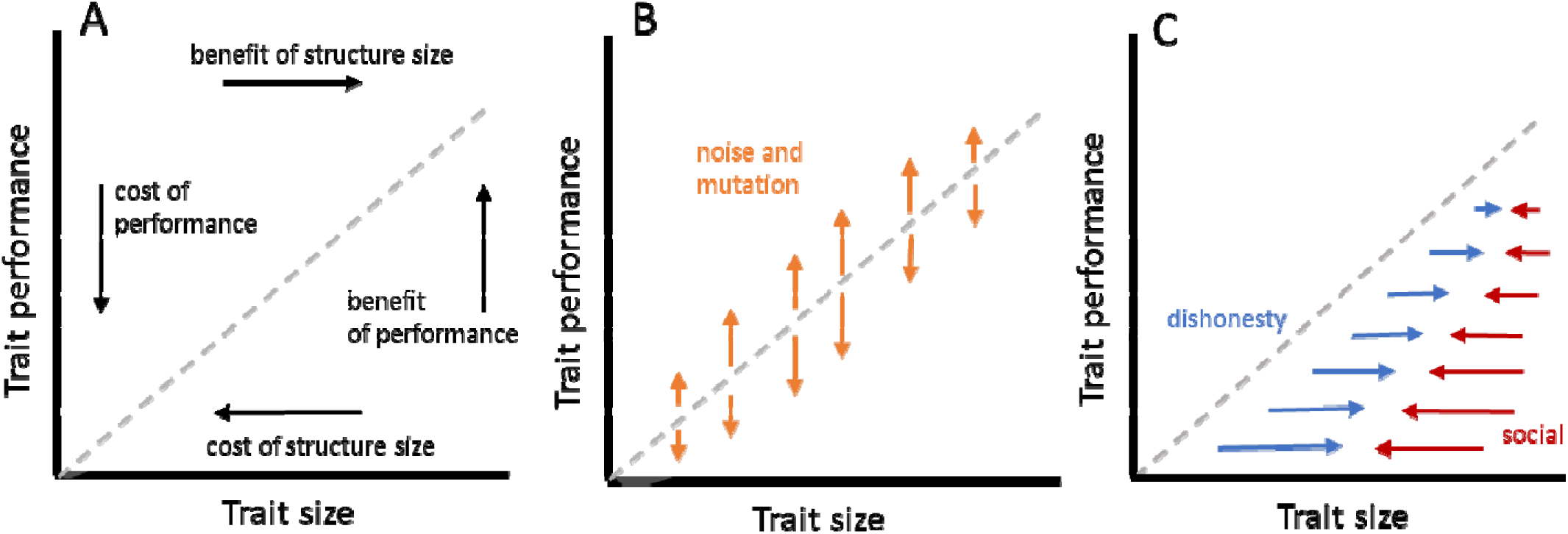
Graphical depiction of the covariation of trait size and performance with different inputs from selective and nonselective pressures. In all scenarios, the gray reference line depicts equal allocation of trait performance and trait size and the gray shaded area represents the morphological space that is being occupied. A) The covariation between trait size and performance in which there is a predictable, reliable relationship between trait size and performance. In this scenario, there are costs and benefits for increasing or decreasing a traits size and performance, indicated by the black arrows. B) The same covariation in A) with the addition of developmental noise and mutation (orange arrows), ultimately increasing the covariation between the size and performance of a signal. C) If the trait in question has evolved for signaling functions, there may be selection for an unreliable (dishonest) relationship between trait size in performance (blue arrows), in which there is selection for large but poor performing signals. However, social policing of dishonest individuals (red arrows) is expected to reduce selection for dishonest signalers.

We measured the strength of each claw with a custom-built force transducer, consisting or two metal plates connected to a load cell and an amplifier. When held close to the plates, a crayfish used its claw to pinch the plates together. The amplifier generated a voltage that was transformed linearly to determine pinching force. We estimated maximal strength of each claw from the best of five consecutive pinches; such measurements have been highly repeatable for other species of crayfish (Graham, Padilla-Perez, et al., 2020; Robbie S Wilson et al., 2009). To confirm this repeatability in our study, we re-measured maximal claw strength on two consecutive days for a subset of individuals of each species.

### Staged territorial encounters

A subset of the males of each non-burrowing species were used to stage encounters between competitors. Burrowing crayfish were not used during these trials because they neither behave aggressively nor defend territories (ref). Prior to staged encounters, we randomly selected half of the crayfish to be focal individuals. Each focal individual experienced in 2-4 staged encounters. Opponents were assigned randomly with the constraint that no opponent could be encountered more than once. At least 24 hours elapsed between the consecutive encounters of each focal individual (ref).

In total, we staged 150 encounters: 40 encounters between males of *C. robustus*, 50 encounters between males of *C. carinirostris*, and 60 encounters between males of *F. obscurus*. Encounters occurred in a glass aquarium (35 × 20 × 15 cm) filled with dechlorinated water and divided by a removable screen. The screen prevented physical contact between crayfish without preventing visual or chemical signaling. We placed a pair of crayfish in the aquarium simultaneously and removed the divider after 5 minutes. Upon removing the divider, crayfish typically engaged in aggression within 5 minutes. Aggression typically consisted of multiple bouts of interlocking claws and grappling, with occasional pinching. Once aggression ensued, we observed the interaction until the winner chased and the loser fled (Wilson et al., 2007). Upon determining a winner, we stopped the trial to prevent injuries. Importantly, some contests were settled solely by displays of claws (refs). Thus, we categorized each encounter according to its final stage of escalation: 1) encounters that ended without physical contact, and 2) encounters that ended after physical contact (e.g., tapping, grappling, or pinching with claws). For our analyses, we considered any encounter that escalated to physical contact as a fight. Collecting data on the final stage of aggression enabled us to disentangle factors promoting dominance prior to fighting and factors promoting dominance during fighting.

### Statistical Analyses

When analyzing the repeatability of claw strength, we conducted a generalized linear mixed model to determine the relationship between maximal strength recordings across consecutive days. Importantly, because each crayfish has two claws, we wanted to control for individual variation. Thus, we used individual crayfish as a random factor.

To determines how each difference species’ claw strength varied within a given size, we regressed the maximal strength of each claw onto the size of each claw (using PC scores). Like previous studies (ref), we compared the fit of models in which the error was ether an exponential, power, constant power, or contestant (ref). To do so, we used the *nmle* library in R (ref) to fit linear models of claw size and claw strength. The most likely model based on the corrected Akaike information criterion (AICc) was used to calculate the residual strengths of each claw.

Having estimated residual claw strengths, we then fit models with maximal claw strength as the dependent variable, and with an interaction between claw size, lifestyle, and sex as independent variables. Furthermore, we included a random intercept associated with individual crayfish. To test the hypothesis that the burrowing lifestyle would lead to reduced selection for claws with immense variation in strength, we compared models with and without lifestyle as a fixed factor. When testing our hypothesis about how sex, species, and lifestyle influence the variation in crayfish claw strength for a given size, we estimated the variance of claw strength for each combination of sex and lifestyle, to see whether adding these parameters provided a better fit to the data than a single variance for all groups. Based on our hypothesis, we expected that the most likely model would be one with a larger variance of claw strength for non-burrowing crawfish than for burrowing crayfish. We also predicted a greater variance in claw for males than for females

When testing our hypothesis about the traits that determine the outcome of territorial aggression in the non-burrowing species, we modeled the factors influencing the probability of the focal crayfish engaging in aggression and the probability of winning a fight. We fit generalized linear mixed models to our data using the lme4 library of the R Statistical Package (Bates et al. 2015). In each case, the dependent variable was coded as a discrete outcome (0 or 1); therefore, we used a model with a binomial distribution of error. To control for the effects of collinearity in our fixed effects, we initially ran separate models of carapace size, claw size, and claw strength difference to see how these effects influenced out independent variables (probability of fighting or probability of winning). Additionally, a random slope of species was added to these models. The single term model with the lowest AIC value was then used for consequent analyses, described below. After determining the best model from the models comparing our fixed effects, we used multimodel averaging to estimate the most likely value of means. Then, we used the MuMIn library (Bartoń, 2013) to fit all possible models to the data.

After fitting all possible models, we calculated the Akaike information criterion and Akaike weight of each model, the latter variable being the probability that the model best describes the data. Finally, we calculated the weighted average of each parameter including estimates from all models. The resulting values of parameters were used to calculate the most likely mean for each combination of factors. This approach eliminates the need to interpret *p* values because all models (including the null model) contributed to the most likely value of each mean.

## RESULTS

### Repeatability of claw strength

The relationship between maximal force produced on consecutive days demonstrates that our measurement of claw force is reliable (Figure X). Regardless, of sex, species, or lifestyle, the claws maximal forces produce on day one of measurements strongly correlate to the maximal forces produced on the second day, demonstrated by the high *r*^2^ values for each species. (Figure X; *C. carinirostris*: *r*^2^ = 0.89, *t*_84_ = 25.71, *p* < 0.001; *C. robsutus*: *r*^2^ = 0.89, *t*_42_ = 18.75, *p* < 0.001; *F. obscurus*: *r*^2^ = 0.91, *t*_37_ = 19.63, *p* < 0.001; *C. monongalensis*: *r*^2^ = 0.74, *t*_32_ = 9.61, *p* < 0.001; *C. dubius*: *r*^2^ = 0.95, *t*_31_ = 24.13, *p* < 0.001; *L. thomai*: *r*^2^ = 0.91, *t*_23_ = 14.89, *p* < 0.001).

**Figure 2.**
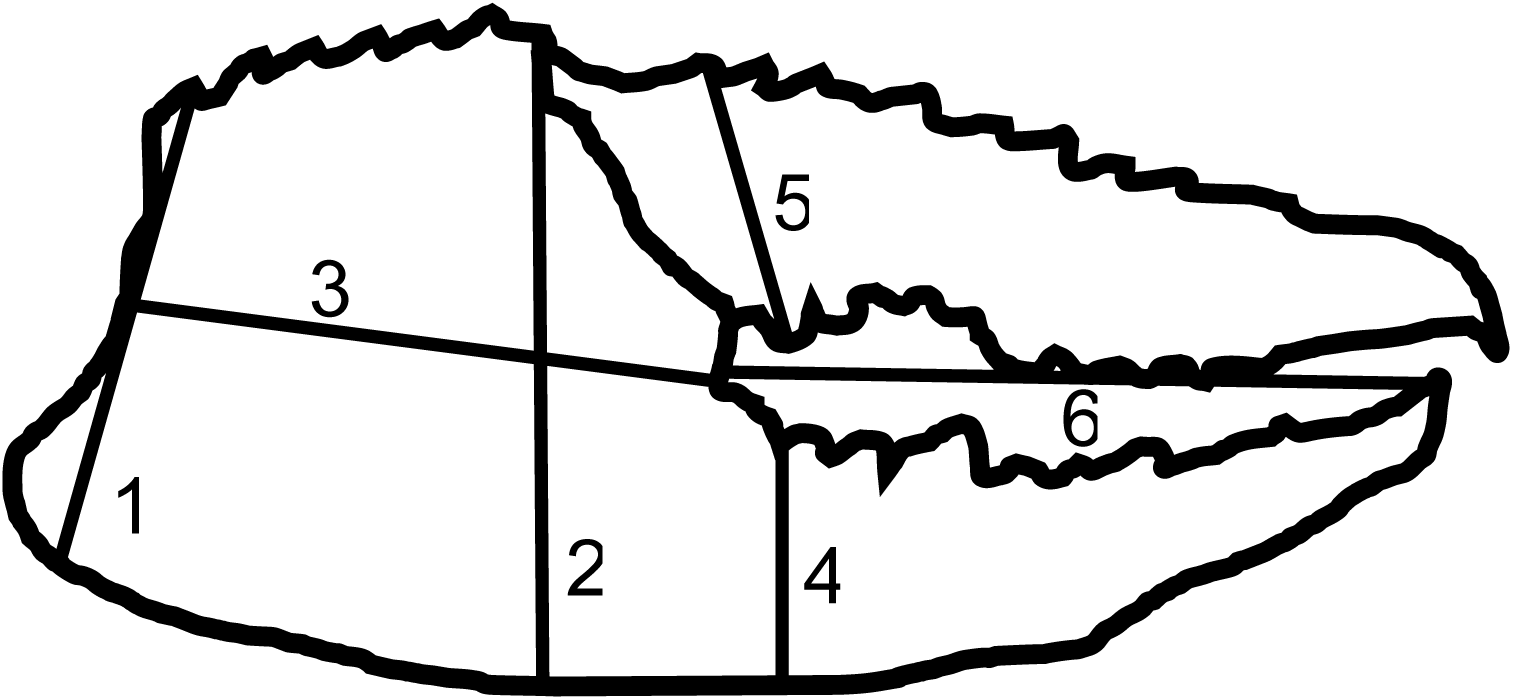
Diagram showing the six linear measurements to describe the size of each claw of the virile crayfish. The six measurements were as follows: 1) width at heel, 2) width at dactyl/manus joint, 3) length of manus from heel to joint, 4) width of pollex at dactyl joint, 5) width of dactyl, 6) and length of pollex from tip to joint.

### Variation in strength among sex, species, and lifestyle

Each species had significant variation in claw strength for a given size, represented by the low *r*^2^ values for the males and females of each sex. (Figure 3; male *C. carinirostris*: *r*^2^ = 0.26, *t*_88_ = 5.592, *p* < 0.001; female *C. Carinirostris*: *r*^2^ = 0.13, *t*_89_ = 3.678, *p* < 0.001; male *C. robustus*: *r*^2^ = 0.23, *t*_56_ = 4.101, *p* < 0.001; female *C. robustus*: *r*^2^ = 0.15, *t*_58_ = 3.278, *p* < 0.002; male *F. obscurus*: *r*^2^ = 0.36, *t*_72_ = 5.158, *p* < 0.001; female *F. obscurus*: *r*^2^ = 0.30, *t*_58_ = 5.055, *p* < 0.001; male *C. monongalensis: r*^2^ = 0.06, *t*_24_ = 1.291, *p* = 0.209; female *C. monongalensis*: *r*^2^ = 0.26, *t*_43_ = 3.904, *p* < 0.001; male *C. dubius*: *r*^2^ = 0.001, *t*_24_ = 0.166, *p* = 0.869; female C. *dubius*: *r*^2^ = 0.27, *t*_36_ = 3.631, *p* < 0.001; male *L. thomai*: *r*^2^ = 0.08, *t*_14_ = 1.14, *p* = 0.273; female *L. thomai*: *r*^2^ = 0.19, *t*_30_ = 2.626, *p* = 0.014). When determining the influence that sex, species, ad lifestyle had on the variation in claw size and claw strength, we found that the lifestyle and sex did influence this variation, but not in the way we predicted. Overall, the variation in the residual strength of both male and female burrowing species (male: SD = 1.372; female: SD = 1.313) was greater than the residual strength variation found in male and female nonburrowing species (male: SD = 1.000; female: SD = 1.217).

**Figure 3.**
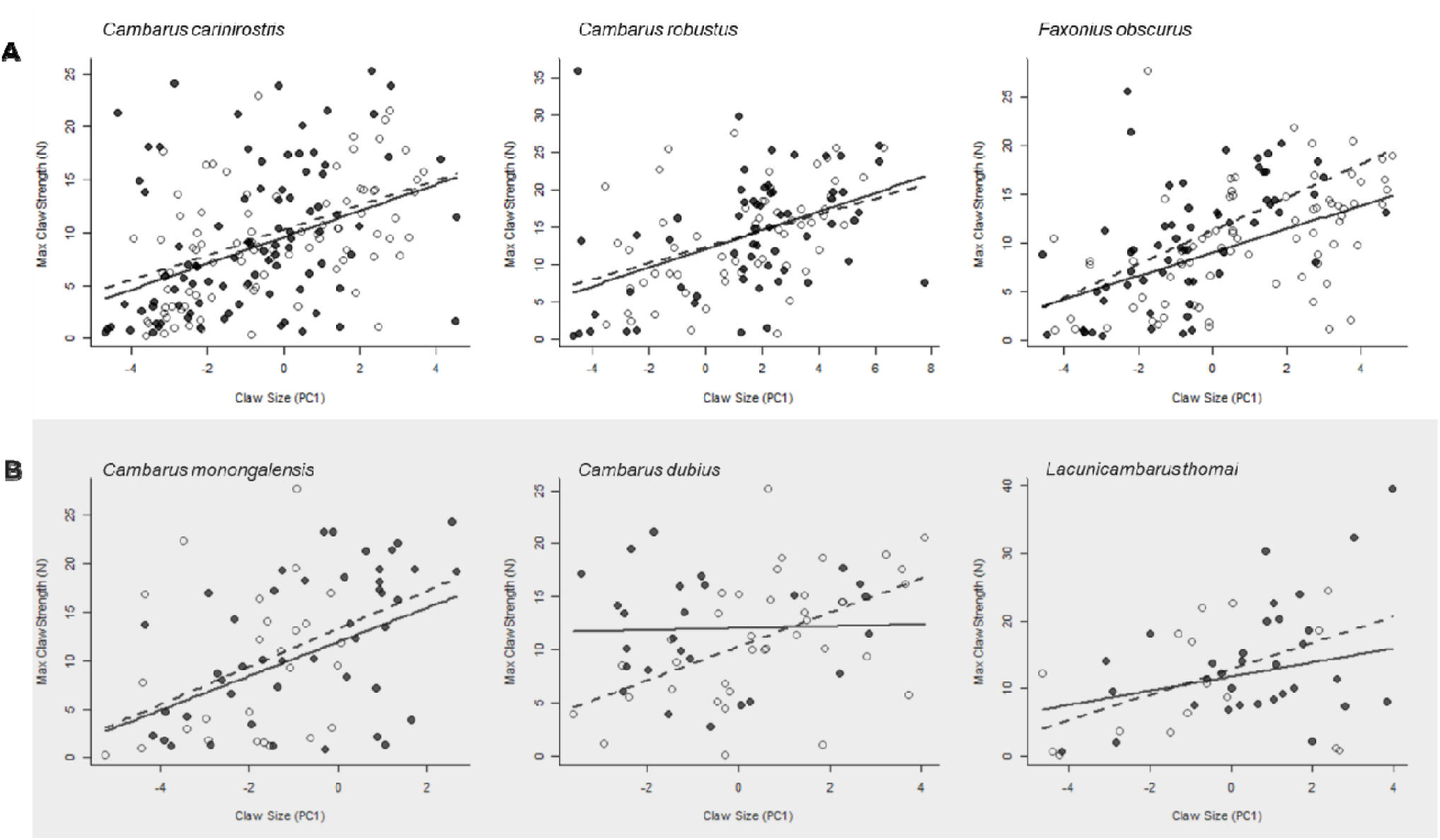
Relation between claw size (PC1) and maximal claw strength in three (**A**) non-burrowing and the (**B**) three burrowing species included in our study. Filled and unfilled circles represent data for males and females, respectively. Furthermore, the full black line and dotted black line represent the linear relationship between males and females, respectively.

### Staged territorial encounters

During staged encounters in which did not escalate to physical aggression, the outcome of these contests was best predicted by the relative body size of the contestants, and not claw size, as we predicted (Fig. 6, Table S2). Furthermore, when we analyzed the outcome of contests that resulted in direct physical fighting, we again found that relative body size best predicted the outcome of these fights, and not the size or strength of claws (Fig. 6, Table S2). In both analyses, the species that were fighting had a small effect, which likely demonstrates subtle differences in the assessment and fighting strategies of each species. (Fig. 6, Table S5).

**Figure 4.**
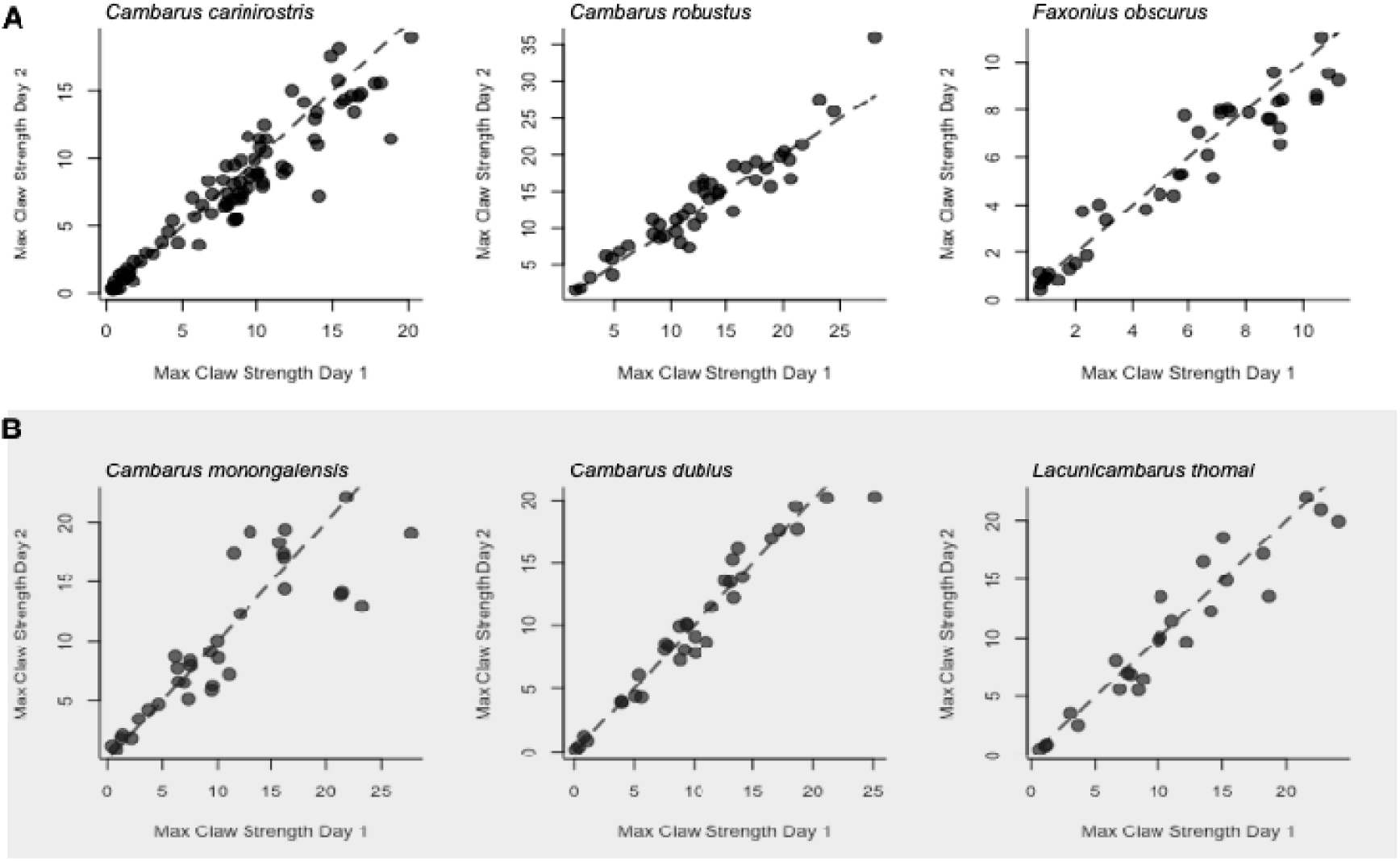
Relation between the strength of the claw in males and females of (**A**) the three non-burrowing species and (**B**) the three burrowing species on day 1 and day 2 controlling for crayfish identity as a random factor in a generalized linear mixed model. The dashed line serves as a reference for the equality of strength between days.

**Figure 5.**
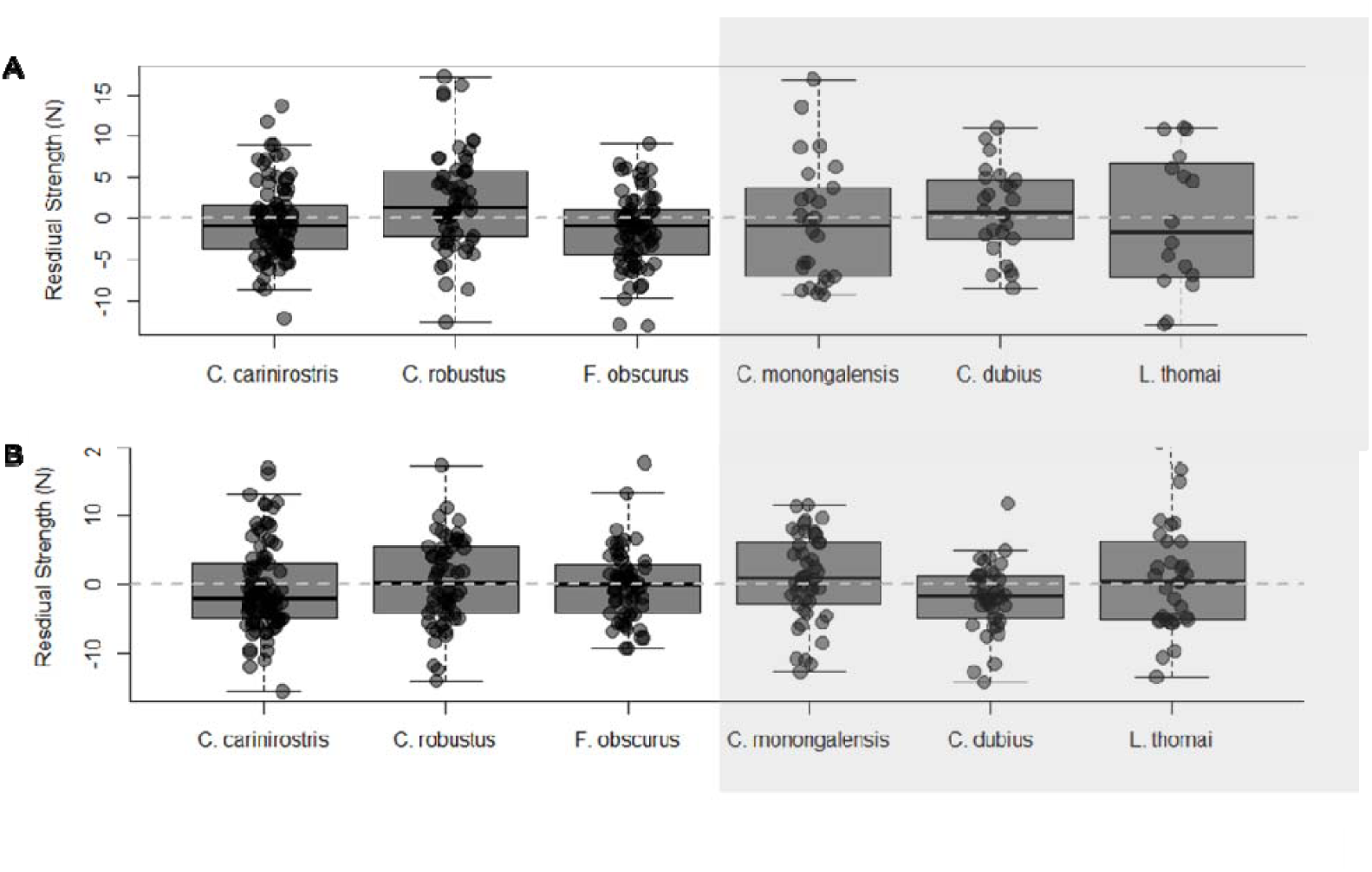
The residual maximal claw strength for (**A**) male and (**B**) females of the six species included in our study. Species within the gray shaded area are the burrowing species, whereas the nonshaded areas represent the non-burrowing species. Contrary to our prediction, the residua variation in claw strength was greater in burrowing species. Error bars represent 95% confidence intervals around the mean residual strength for each species. The dashed references line depicts a residual strength value of zero.

**Figure 6.**
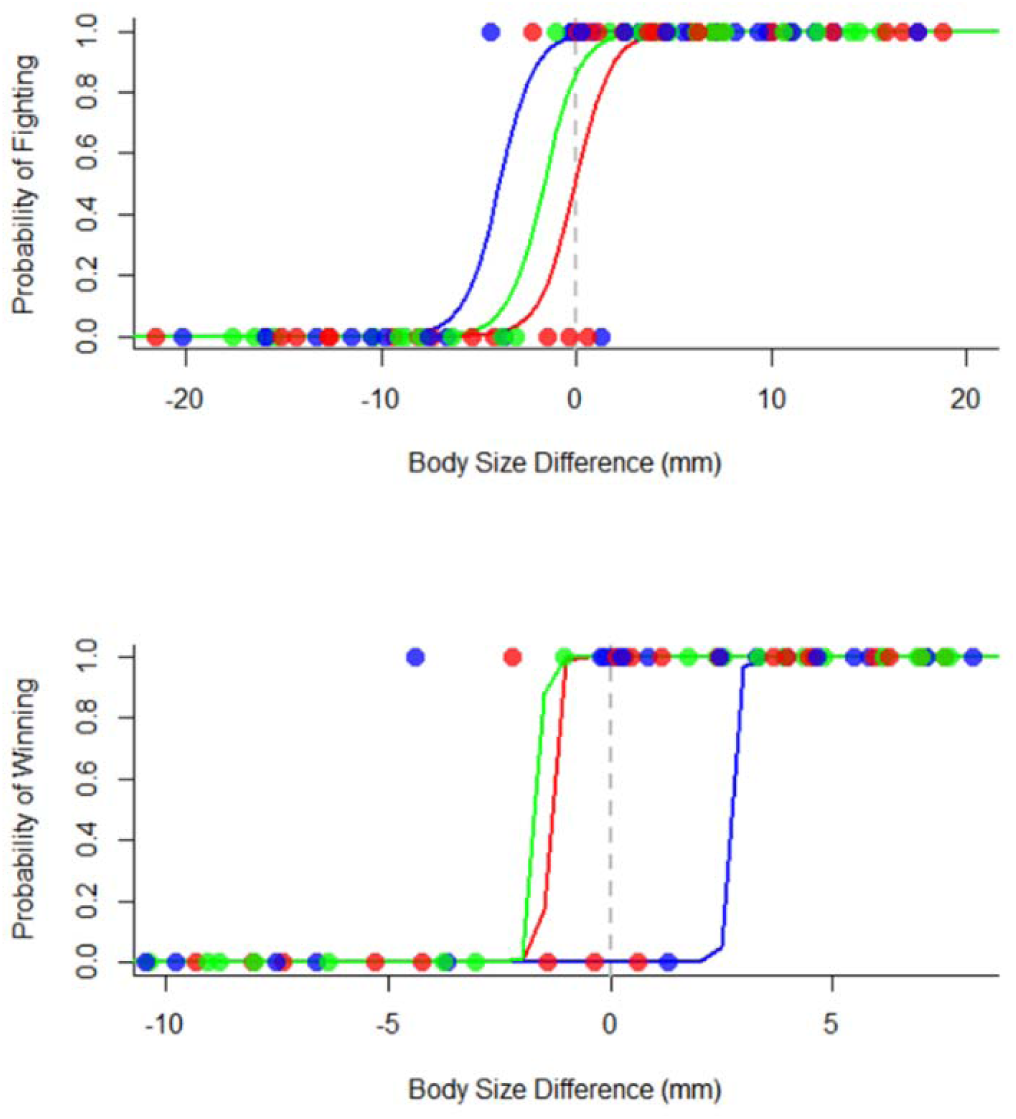
(**A**) Probability of escalating aggression to physical combat based on the difference in claw strength between opponents. (**B**) Probability of winning a fight based on the difference in carapace length between opponents. Circles indicate the outcome of each encounter (0 or 1, where 1 represents winning). The lines represent the fitted values of statistical model that describes our data. In both graphs, the red dots and lines represent the data from *C. carinistostris*. Blue dots and lines represent data *C. robstus*, and green dots and lines repsent data from *F. obscurus* aggression. The dotted line provides a reference for zero difference in body size.

## DISCUSSION

Despite substantial effort to study dishonest signaling in natural systems; it is difficult to classify whether a signal is honest or dishonest. In fact, our ability to study dishonest signals is entirely reliant on our ability to detect and describe the relative honesty of such signals. Initially, dishonest signals were described subjectively and qualitatively as when a sender, but not receiver benefitted from communication. Further, signals were also determined to be dishonest if there was a breakdown between the signals design and the information being communicated (Adams & Caldwell, 1990; Bradbury & Vehrencamp, 1998; Maynard Smith, 1982; Searcy & Nowicki, 2005). For example, alarm calls of birds signal to receivers that a threat is near; thus communicating honest information. But in some scenarios, birds will emit dishonest alarm calls (Flower, 2011; Flower, Gribble, & Ridley, 2014; Munn, 1986; Møller; 1988). When a dishonest alarm call is signaled, receivers still respond as if a threat is present, which gives the signaler an opportunity to forage or exploit potential resources with little competition from the fleeing receivers. Thus, the breakdown of the relationship between the alarm call (the signal) and the behavior of receivers (hiding as if the alarm call were true) can be used to determine the honesty or dishonesty of the single. Although these descriptions of dishonesty were paramount in our early studies, such descriptive and informal definitions of dishonesty made it difficult to ask ecological and evolutionary relevant questions as to how and why such signals can persist.

More recently, quantitative methods of studying the honesty of animal signals have bolstered our understanding dishonest signaling. Specially, the use of correlation coefficients and signal residuals have enabled researchers to statistically characterize and compare the relative reliability of a signal within and between animal species (Akçay, Campbell, & Beecher, 2014; Briffa, 2006; Carazo & Font, 2010, 2013; Hughes, 2000). For example, by calculating the correlation between the intensity of begging calls and the condition of chicks, Caro & Colleagues were able to investigate how the degree of sibling conflict influenced the relative honesty or dishonesty of begging calls across 60 species of birds (Caro, West, & Griffin, 2016). Species with tighter correlations between signal intensity and condition communicate honesty; whereas other species exhibit little correlation between intensity and condition – communicating dishonestly. The honesty of signals is therefore best understood as a statistical property of the system. In a similar way, by studying the residual variation from the relationship of the size or magnitude of the signal and the quality or intention being signaled, researchers can analyze the behavior of honest (positive signal residuals) or dishonest (negative signal residuals) signals effect on signaling systems (Briffa, 2006; Hughes, 2000; R. S. Wilson & Angilletta, 2015). Interestingly, in scenarios where the interest of signalers and receivers are opposed, such as territorial aggression between crustaceans, the use of dishonest signals seem to be more are prevalent (R. S. Wilson & Angilletta, 2015). However, more recent studies in crayfish have demonstrated that despite a weak correlation between the size and strength of male crayfish claws, that the claws did not function as unreliable signals (Graham, Padilla-Perez, et al., 2020).

The results of our study infringe on the validity of the traditional methods of determining the honesty of animal signals. Despite predicting that there should be reduced variation in the burrowing crayfish claws that do not function as signals, we found similar variation in the sizes and strength of their claws when compared to non-burrowing species. Therefore, inherent noise within this system could cloud biologist’s interpretation of the relative honesty or dishonest of such signals (Carazo & Font, 2010, 2013). Although variation in the size and quality of animal signals have been proposed as evidence of dishonesty, our study demonstrates that just being a putative signals unreliably predicts quality, this is not evidence of dishonesty. In staged interactions between non-burrowing species, we found that despite being extremely variable; their claws are unlikely to function as signals because claw size did not determine whether or not individuals engaged in combat. Interestingly, although the claws of many other crustaceans such as fiddler crab’s species have been heavily studied for their function as dishonest signals (Bywater, Seebacher, & Wilson, 2015; Bywater & Wilson, 2012; R. S. Wilson & Angilletta, 2015), our finds support for unreliable, but not dishonest signaling in crayfish. Further, although previous studies have suggested that dishonest signaling should be more common in male crayfish, we found equal rates of unreliability between male and female claws.

We propose several explanations as to why the claws of non-burrowing and burrowing crayfish species exhibit have similar degrees variation. First, the claws of the nonburrowing crayfish in our study may have been selected for their function purely as weapons, with little to no signaling functions (Mccullough, Miller, & Emlen, 2016). In this way, there does not necessarily need to be selection for increased variation in the size and performance of non-burrowing species claws. Because the results of our staged interactions indicated that the non-burrowing species we studied were unlikely to function as signal’s, there may be selected to reduce variation in the size and performance of non-burrowers claws in order for them to operate as efficient weapons. Such size-based limitations are common in animal weapons that are lever systems, like crayfish claws or the hindlegs of leaf-footed bugs (Dennenmoser & Christy, 2013; Levinton & Allen, 2005; O’Brien & Boisseau, 2018; Palaoro, Peixoto, Benso-Lopes, Boligon, & Santos, 2020). Secondly, despite differences in the function of burrowing and non-burrower claws, these claws could share a similar noise inducing mechanism producing the degree of variation we observed. Because crustaceans such as crayfish require to go through a molting cycle in order to grow, they are in a constant flux of muscular and cuticle regeneration (Chang, 1995). Interestingly, the large muscle inside crustaceans’ claws must undergo significant muscular atrophy in order to molt (West, 1997). Therefore, in mature crayfish, which molt one or multiple times a year, they must go through significant changes in musculature during this time. Although the exoskeleton of crayfish hardens fairly quickly after a molt, regeneration of the atrophied claw muscles may take longer to full regrow. Because the strength of crustacean claws is likely determined by the condition of the individual during a molting cycle; there may be significant variation in the strength that can be produced by the claw. Therefore, although the function’s of burrowing and non-burrowing claws may be separate, both may face substantial variation in the strength of their claws within and between molting cycles; producing the observable patterns of variation we found. In some species of crayfish that have dishonest signals, such as *Cherax dispar*, this may explain the variation found in their claws to function as dishonest signals. In other species, such as the ones we studied, this variation may be too strong to select for dishonesty; ultimately making the system too unreliable for dishonest signals to be maintained.

Throughout our study, we have highlighted the difficulties of identifying and studying dishonest signal’s in natural system of communication. We suggest that the ideal scenario to determine the honesty of signal’s is through signal manipulation studies. By manipulating the size, color, or intensity of a signal, researchers have been able to test the function of such signals in different contexts. Such signal manipulation studies have been lucrative in our understanding of the maintenance of conventional signals; which have no direct link to individual quality. Experimental increase or decrease in the size or color of a putative signal allows explicit tests of the putative signals function. Therefore, by manipulating the size of the signal, researchers can experimentally create individuals with disproportionately larger or small signals. In crayfish and other weapon bearing species, it may be possible to artificial increase or decrease the size of their weapons to test the function of the signal. For example, by using the molted claw of a larger crayfish, researchers could attach an artificially large claw atop of a smaller clawed individual; effected increasing the size of the claw. If the claw functions as a signal, this larger claw should change the size of opponents willing to engage in aggression. Thus, such manipulation would enable tests of the effect of claw size as signals through alterations of the covariation between the size of a signal and the quality being signaled. Therefore, despite the inherent noise within our system, our study cautions future interpretation’s as variation as dishonesty and furthermore, more explicit tests of signaling must be done before determining the function of a structure as a signal.

