## Supplemental Materials for "Separating noise and function in systems of animal communication: a comparative study of aggressive signaling in crayfish"

**Table S1.** Principal components loadings for claw size measurements of crayfish. The values presented are relative contributions of the six claw measurements to the variation explained by each principal component. PC1 values all having loadings in the same direction and best explain the variation in overall claw size.

| Claw Measurement |  | PC1 | PC2 | PC3 |
| --- | --- | --- | --- | --- |
| 1 | Width at heel | 0.411 | 0.312 | 0.134 |
| 2 | Width at dactyl/manus joint | 0.418 | 0.158 | 0.012 |
| 3 | Length of manus from heel to joint | 0.403 | 0.515 | -0.518 |
| 4 | Width of pollex at dactyl joint | 0.407 | -0.414 | 0.429 |
| 5 | Width of dactyl | 0.413 | 0.069 | 0.470 |
| 6 | Length of pollex from tip to joint | 0.396 | -0.651 | -0.556 |
| Eigenvalue |  | 0.729 | -0.542 | -0.135 |
| % of variance |  | 93.92 | 3.45 | 1.11 |
| Cumulative |  | 93.92 | 97.37 | 98.48 |

**Table S2.** The most likely models predicting the probability of engaging in aggression

| **Model** | ***df*** | **Log likelihood** | **AICc** | **ΔAICc** | ***w*** |
| --- | --- | --- | --- | --- | --- |
| body size difference | 3 | -12.87 | 32.00 | 0.00 | 0.65 |
| body size difference + species | 5 | -11.27 | 33.19 | 1.20 | 0.35 |
| null | 2 | -50.57 | 105.27 | 73.28 | 0.00 |
| species | 4 | -50.38 | 109.19 | 77.19 | 0.00 |

All models contained an intercept and error terms associated with the identity of the focal crayfish. For each model, the degrees of freedom (*df*), the corrected Akaike information criterion (AICc), Akaike weight (*w*) and the log likelihood are reported. Models were ranked according to their corrected AICc.

**Table S3.** Coefficients and standard errors for the model of probability of winning a fight with an opponent crayfish, based on full model averaging.

| *Independent variable* | *Coefficient* | *SE* |
| --- | --- | --- |
| intercept | 1.2475 | 1.1971 |
| body size difference | 1.0331 | 0.3196 |
| *C. robustus* | 4.2587 | 3.1332 |
| *F. obscurus* | 1.7607 | 1.9610 |

**Table S4.** The most likely models predicting the probability of winning a fight

| **Model** | ***df*** |  | **Log likelihood** | **AICc** | **ΔAICc** | ***w*** |
| --- | --- | --- | --- | --- | --- | --- |
| body size difference + species | 5 |  | --12.65 | 36.84 | 0.00 | 0.50 |
| body size difference | 3 |  | -15.69 | 37.97 | 1.13 | 0.28 |
| body size difference + claw size difference | 4 |  | -14.91 | 38.83 | 1.99 | 0.18 |
| body size difference + claw size difference + species | 6 |  | -14.01 | 42.24 | 5.40 | 0.03 |
| claw size difference | 3 |  | -22.21 | 51.00 | 14.17 | 0.00 |
| claw size difference + species | 5 |  | -21.38 | 54.30 | 17.46 | 0.00 |
| null | 2 |  | -31.17 | 66.62 | 29.79 | 0.00 |
| species | 4 |  | -29.74 | 68.49 | 31.65 | 0.00 |

All models contained an intercept and error terms associated with the identity of the focal crayfish. For each model, the degrees of freedom (*df*), the corrected Akaike information criterion (AICc), Akaike weight (*w*) and the log likelihood are reported. Models were ranked according to their corrected AICc.

**Table S5.** Coefficients and standard errors for the model of probability of winning a fight with an opponent crayfish, based on full model averaging.

| *Independent variable* | *Coefficient* | *SE* |
| --- | --- | --- |
| intercept | 8.1567 | 10.7999 |
| body size difference | 7.7786 | 5.1074 |
| *C. robustus* | -25.8976 | 26.6746 |
| *F. obscurus* | 1.3855 | 6.9467 |
| claw size difference | 0.6765 | 2.0474 |

**Figure S1.** Scatter plot of the first two measurements (PC1 and PC2) from the Principal Component Analysis of crayfish claw size. Claw sizes are variable across species, demonstrated by the clustering of individual's species claws. Circles and triangles represent individual claws from male and female species, respectively. Species are represented by different colors.

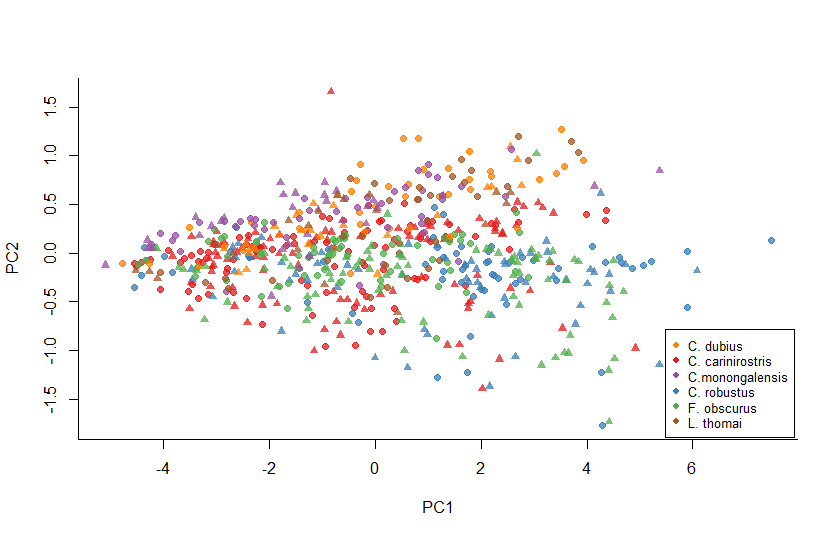
